# Cogno-Vest: A Torso-Worn, Force Display to Experimentally Induce Specific Hallucinations and Related Bodily Sensations

**DOI:** 10.1101/2020.06.23.167551

**Authors:** Atena Fadaei J., Kenny Jeanmonod, Olivier A. Kannape, Jevita Potheegadoo, Hannes Bleuler, Masayuki Hara, Olaf Blanke

**Author notes:** These two authors contributed equally to this work (Masayuki Hara, Olaf Blanke).

## Abstract

Recent advances in virtual reality and robotic technologies have allowed researchers to explore the mechanisms underlying bodily aspects of self-consciousness which are largely attributed to the multisensory and sensorimotor processing of bodily signals (bodily self-consciousness, BSC). One key contribution to BSC, that is currently poorly addressed due to the lack of a wearable solution, concerns realistic collision sensations on the torso. Here, we introduce and validate a novel torso-worn force display, the Cogno-vest, to provide mechanical touch on the user’s back in a sensorimotor perception experiment. In a first empirical study, we characterized human finger poking (N=28). In order to match these poking characteristics and meet the wearability criteria, we used bi-directional, push-pull solenoids as a force actuator in the Cogno-vest. Subsequently, and based on an iterative, multidisciplinary design procedure, a body-conforming, unisex, torso-worn force display was prototyped. Finally, we conducted a behavioral study that investigated BSC in 25 healthy participants by introducing conflicting sensorimotor signals between their hand and torso (back). Using the final reiteration of the Cogno-vest we successfully replicated previous findings on illusory states of BSC, characterized by presence hallucinations (PH) and passivity symptoms, and achieved higher illusion ratings compared to static conditions used in prior studies.

## I. Introduction

Wearable haptic displays for the torso have been gaining importance in recent years in a variety of applications, ranging from virtual environments [1], [2] to navigation [3]–[9], affective stimulation [10]–[16], rehabilitation [17], [18] as well as sensory-substitution [15], [19]–[21]. Torso-worn haptic displays range from precise, focal applications to broad surfaces, and convey haptic information without competing for audio-visual attentional resources, while offering a hands-free solution [22]. Torso-worn tactile stimulators can easily be used in a portable, wearable arrangement (e.g., in the form of an actuator vest), and the information provided is mostly processed automatically, imposing relatively low attentional loading on the user [4].

Preliminary works in the field of torso-based haptic displays focused on navigation and spatial orientation purposes as the human torso is often referred to as the location and central reference frame for the human body [23]. In two seminal studies, Rupert et al. developed and tested different vibrotactile torso-worn displays to facilitate orientation awareness for pilots in unusual acceleration environments [4], [24]. Van Erp et al. deployed a torso-worn vibrotactile display as a pedestrian navigation system [25]. They also carried out a series of systematic experiments to determine vibrotactile perception on the torso in humans. Several studies have been published on the use of torso-based haptic displays, as a tactile-visual sensory substitution system, to aid visually impaired people. The VibroVision vest, developed by Wacker et al. [19], comprised an array of 16 x 8 vibrators and could convert visuospatial information into a 2D vibration image on the abdomen. Similarly, the Tactile Vision Substitution (TVS) system was designed to capture visual information from the surrounding environment using a video camera and to deliver feedback to the skin of the back, abdomen, or thigh via an array of actuators [15].

In addition, previous studies have demonstrated that vibrotactile actuators are able to transmit physical information either as a simple cue to indicate contact location in virtual environments or a complex one to convey an object’s physical properties. For instance, Lindeman et al. designed TactaVest [26], a torso-based vibrotactile interface, to deliver the users feedback about collisions with virtual objects in military simulation. Previous studies have also demonstrated the application of torso-based tactile displays for the communication of affective touch. Lemmens et al. developed a wearable vibrotactile jacket, involving 64 vibrators, intended to intensify the emotional immersion experience while watching movies [27]. Arafsha et al. designed the Emojacket for enhancing immersion while watching movies and gaming experience, containing a combination of vibrotactile and heat actuators to display several universal emotions, along with several emotional reactions such as a hug, poke, tickle, or touch [10]. Lentini et al. designed a tactile gesture authoring system able to translate hand gestures into vibrotactile stimuli rendered through a haptic jacket [11].

Researchers in cognitive neuroscience have recently highlighted the crucial role of the processing of somatosensory signals from the torso for global aspects of Bodily Self Consciousness (BSC), a crucial brain mechanism of self-consciousness based on perceptual mechanisms associated with the integration of multisensory and sensorimotor bodily signals [28], [29]. In these studies, participants were exposed to a large variety of a different conflicting combinations (spatial and/or temporal mismatch) of torso-based touch feedback (e.g., stroking/tapping on the chest/back) and other sensorimotor bodily signals (e.g., visual stimuli and motor action). As the conflicting spatio-temporal feedback prevents normal multisensory integration of bodily signals, participants experienced different illusory own-body perceptions, affecting both self-location and self-identification.

So far, torso-worn haptic displays have rarely been used for investigating global BSC. Ehrsson [18] and Lenggenhager [30] studied out-of-body and full-body illusions by providing participants with an image of their filmed body or a virtual body on a Head Mounted Display (HMD) while an experimenter stroked their torso with a stick. As a result of multisensory stimulation, participants reported that they are located outside their physical bodies, self-identified with the virtual body, and had the impression of looking at themselves from this perspective. A significant limitation of such manual touch-feedback is the inherent variability in both-touch location and timing, and its effects on replicability.

Recently, Blanke et al. used a robotic system to apply sensorimotor manipulations between the hand and torso of blindfolded, sound-isolated participants [31]. Participants were asked to perform poking movements with both hands, by using the front haptic device placed in front of them, while receiving tactile stimuli on their back from another robotic device synchronously or asynchronously with their hand movements. In the synchronous condition, participants reported the experience of touching their own back (illusory self-touch). Interestingly, in the asynchronous condition, they reported a reduction in self-touch sensations as well as the impression that someone else was behind them (presence hallucination, PH) and touching them (passivity experience). It has been argued that such robot-induced experiences are similar to symptomatic PH reported by neurological and psychiatric patients (e.g., Parkinson’s disease and schizophrenia respectively) [32].

Despite recent progress in robotic control for inducing different types of illusory own-body perceptions, the use of such a system is limited to the laboratory environment in which participants remain stationary. Therefore, to investigate the different facets of global BSC, more versatile haptic technology is required that allows us to manipulate tactile stimuli on the torso, together with other sensory modalities, in a complex and dynamic environment. In our previous research [33], we partly replicated the results of [31] using a custom-made torso-worn vibrotactile garment, reporting an effect of synchrony for passivity experiences (“It was as if someone else was touching my back.”), but failed to significantly modulate self-touch (“I felt as if I was touching my back with my finger”) and PH as observed in [31]. We concluded that a simple vibratory stimulus may not be sufficient in substituting collision-type touch stimuli. However, to provide mechanical touches or simulate collisions in virtual environments, it has been shown that force feedback may enhance immersion [34]. To the best of our knowledge, most torso-worn displays described in the literature provide vibrotactile feedback whereas few publications have used force stimulators to the human torso to provide physical interactions in virtual environments. Force jacket was made of pneumatically actuated airbags to provide strong and variable forces to the torso along with vibrotactile sensations [35]. However, this system included a series of air tubes connected to a large air compressor and vacuum resulting in a very bulky setup (neither mobile nor portable) and confining for the users. More recently, Al-Sada et al. [36] designed the HapticSnakes, a snake-like waist-worn robot that can deliver different types of haptic feedback to the upper body via an exchangeable end effector attached to the waist. One shortcoming of their approach is that the end effector movements can be limiting for the user’s hands and body mobility especially in VR environments as they use hand controller while their vision is occluded. Their design also introduced an unavoidable delay since the robotic arm required to move to different points to apply feedback.

A promising solution could be integrating light force actuators into torso-worn garments allowing users to wear it on a regular basis with minimum motion constraints. Yet, due to the large morphological differences in the torso area (within and between-subject variability), forming and fitting the torso-worn haptic display to the user’s body is another crucial design challenge [37]. Design features should therefore preferably enable the actuator positions to be adjustable on the user’s skin. However, previous studies have not addressed this problem adequately and little attention has been paid to the garment design.

Extending our previous work [33], we here address the complexity of designing a body-conforming, torso-worn, haptic display and propose a multidisciplinary approach, including robotics, fashion design, and 3D printing technologies: Cogno-vest is a novel, portable torso-worn force display, developed to provide human-like poking stimuli on the user’s back and investigate robotically-induced altered own-body perception, including PH, by extending the paradigm proposed by Blanke et al. [31] (experiment 2) to a wearable system.

The remainder of the paper is organized as follows: In section II, we describe the design concepts, realization, and validation of Cogno-vest. Section III describes the experimental scenario and results for the own body perception experiment. Section IV discusses the results and examines possible limitations. Finally, section V presents our conclusions.

## II. Novel Wearable setup to induce bodily illusions

A schematic view of the stationary experimental setup, used in study [31] (experiments 2 and 3), is represented in Fig. 1(a). The setup consists of a haptic device (Geomagic Touch, 3D Systems), i.e. the front haptic interface, and a three degree-of-freedom (DOF) robot, i.e. the back robot. The front haptic interface controls the position of the back robot, which causes complete conjunction between the movements of the two robots. Participants manipulated the front haptic interface using their right index finger (via 3D printed finger support) to control the position of the back robot which in turn provides touch-cues to the participant’s back.

**Fig. 1.**
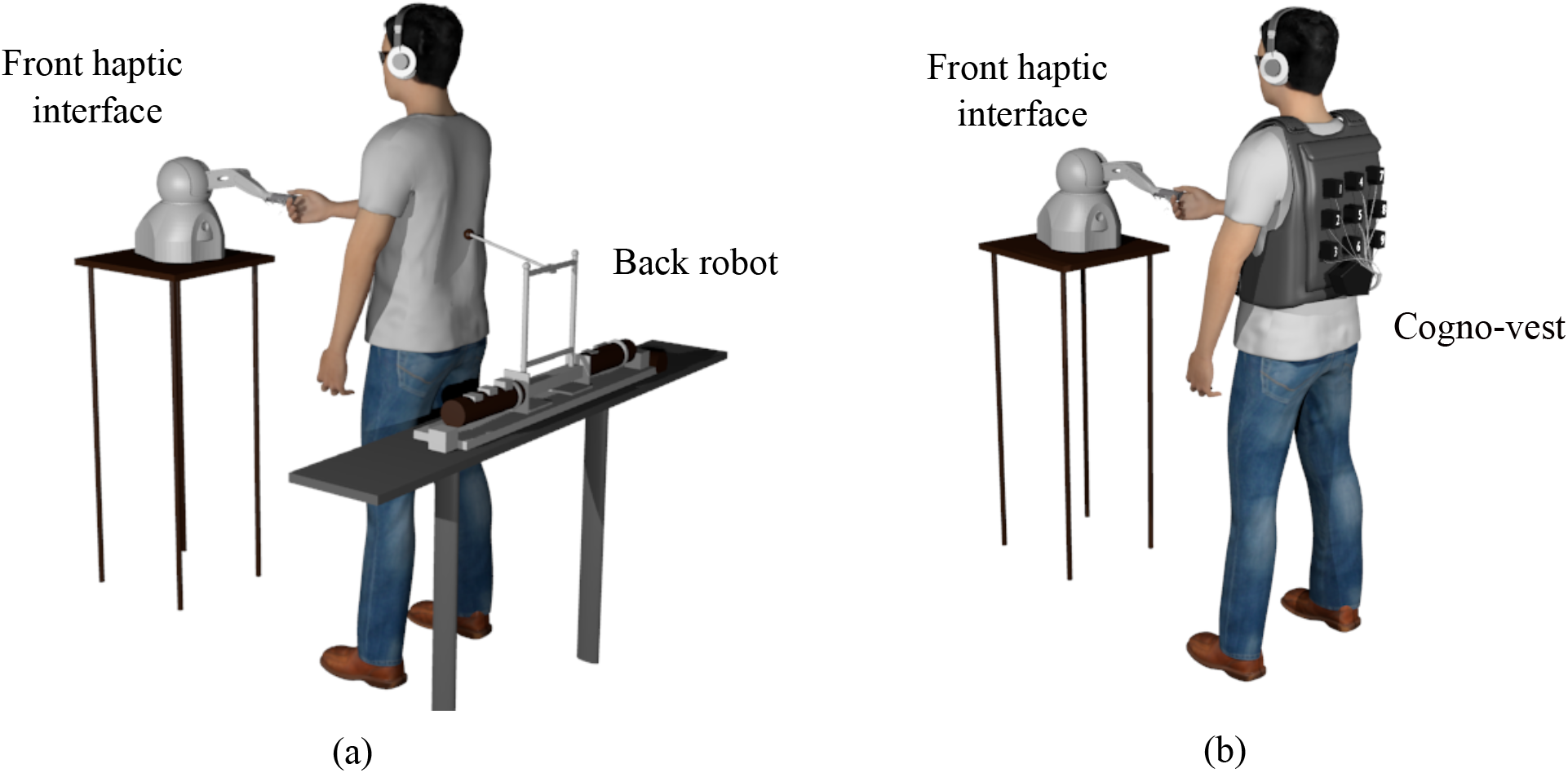
Schematic views of the classical (a) and Cogno-vest setups (b) for experimentally inducing PH and relevant bodily illusions. (a) The setup comprised two robots, the front haptic interface, Geomagic Touch, and a three degree-of-freedom (DOF) follower robot. (b) In the novel setup, the torso-worn force display replaced the robotic arm.

The novel experimental setup is illustrated in Fig. 1(b). In the experimental arrangement, the back robot has been replaced with a torso-worn force display (Fig. 1(b)), called Cogno-vest, to provide the mechanical touch on the participants’ back. The same front haptic interface as in study [31] is used. To assess the spatial correspondence between the continuous hand workspace and discrete back workspace (as a limited number of actuators was employed on the torso-worn display side), the hand exploration area (160 W x 120 H x 120 D mm) was divided into nine square sections, according to the arrangement of the nine on-off actuators embedded in the Cogno-vest. Each square was around 30 x 30 mm while the effective depth (i.e. the depth threshold that, if exceeded, activates one of the actuators, depending on the level of height or width) was considered at the mid-depth. Depending on the participant’s finger poking position in the front robot’s workspace, the corresponding actuator was activated on the back (see Movie S1). The actuator remained operating until particiapnt’s finger departs that actuator’s specific area. In this section, we describe the design concepts, implementation, and characterization of Cogno-vest.

### A. Finger Poking Characterization and Actuator Selection

In order to induce the intended bodily illusions the actuator garment should provide force characteristics comparable to those of human-finger poking, i.e. the act of prodding someone with your index finger, as if to get their attention. Prior to actuator selection, we therefore characterized human finger poking with N=28 participants (14 females, age: 27.6±5.1 years) and quantified peak poking force (*F*_PP_), poking duration (*T*_PD_), and poking interval (*T*_PI_) (see Fig. 2(b)). Fig. 2(a) illustrates the experimental environment. The testbed included a 3D-axis force sensor (OMD-10-SE-10N, OptoForce) attached to a base plate and fixed on a desk in front of the seated participant. Participants were asked to touch the force sensor as if they were poking some one’s back. They were completely free in producing finger poking and no instruction was provided by the experimenter. We gave them tens of second to become familiar with the task, after we recorded data for 1 minute. Fig. 2(c) represents the sample result for one of the participants, while Fig. 2(b) shows the zoom-in in the range of 0-3 s. The results for 28 participants showed that peak poking force and poking duration were *F*_PP_ = 2.15 ± 0.28 N and *T*_PD_ = 0.22±0.03 s respectively. Participants performed finger poking with a time interval of *T*_PI_ = 0.6 ± 0.07 s.

**Fig. 2.**
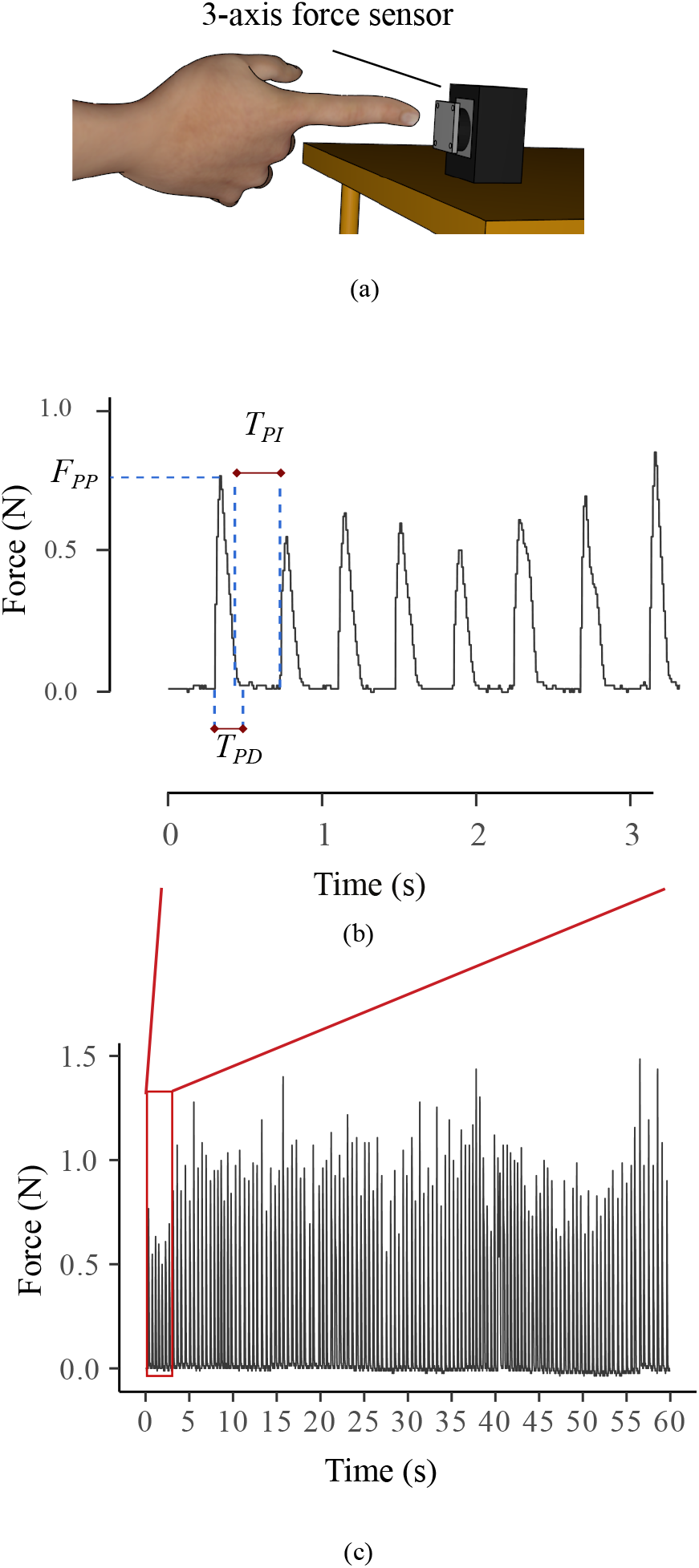
Finger poking characterization experiment’s setup and result. (a) Experimental environment including a 3-D axis force sensor fixed on a the table. (b), (c) Results of finger poking experiment for one of the participants during one minute (c) with special zoom-in in the range of 0-3 s (b). the plot (b) also contains illustrative view of quantitive parameters, used to describe finger poking results (peak poking force (*F*_PP_), poking duration (*T*_PD_), and poking interval (*T*_PD_)).

Based on these results bi-directional, push-pull solenoid actuators that can provide mechanical touch perpendicular to the skin were included in the final design. The solenoid that best matched human figure poking had a starting force of 5 N (12 VDC), a shaft length of 5.5 mm and weighed 39 grams.

### B. Cogno-Vest

#### 1) Electronics and Software Development

In a previous study [33], we reported that an array of 3 by 3 tactile actuators on the back is sufficient to create the sensation that the entire back is covered with actuators. This is due to the poor tactile spatial acuity of the human back [38], and explains the limited number of solenoid actuators used, in our case 9. Solenoids were placed at the center-to-center distance of 60 mm, which approximates the tactile localization threshold of the human back [39], [40]. As a result, the dimension of the back workspace is, approximately, two times larger than that of the hand. Yet, according to our previous study with the vibrator setup and the initial user testing experience with the current setup, the matching ratio of 1:2 between the hand and the back workspace seems natural to participants and, indeed, they are not able to perceive the difference. This observation points to the poor tactile spatial discrimination on the human back.

The Fig. 3(a) presents the system architecture. The actuators are controlled using an Arduino Mega 2560, which connects via a Bluetooth module to a host PC. The controller board can be fully portable (battery-powered) or tethered for more extended studies. The sampling time for the front haptic interface is 1 ms. However, the host PC reads the finger’s position data only every 60 ms, and sends the processed position data in terms of the solenoid activation status (solenoid ID) every 100 ms (this value was adjusted to avoid jittering behavior of solenoids especially at the border of two squares) to the controller board.

**Fig. 3.**
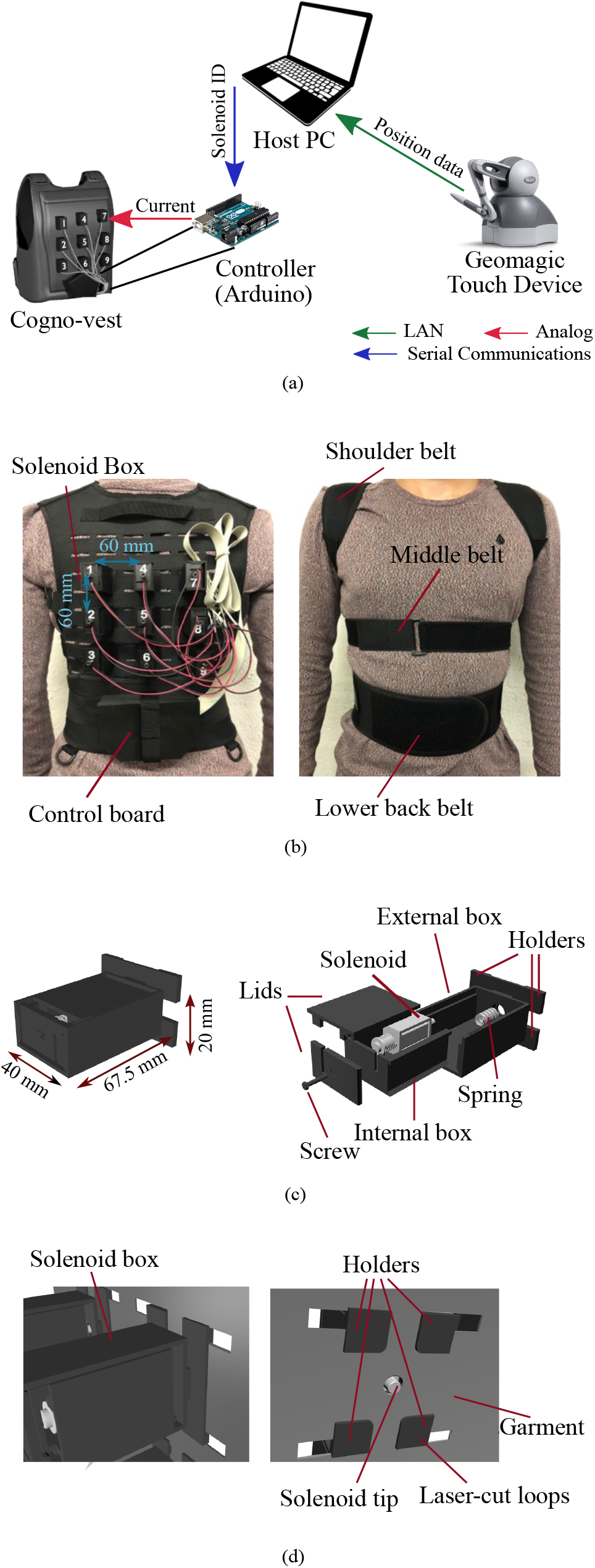
Cogno-vest components. (a) System architecture. (b) Cogno-vest on the participant, back view (left) and front view (right). (c) Mechanically-adjustable box, box dimension (up-left), box components (right). (d) Mounting solenoid boxes on the wearable garment, exterior view (left) and interior view (left).

A customized GUI was implemented in the Qt platform (free and open source platform to create GUI) to provide a convenient interface for controlling haptic stimulations by the solenoids and handling experiments with the Cogno-vest.

#### 2) Garment Design

We designed a new actuator brace for solenoid actuators by considering the following design criteria, suggested by our previous study [33] and literature [26], [37]:

- It should keep actuators snug against the skin, even during walking and movement.
- It should be as light as possible and comfortable to wear, possibly for longer periods of usage.
- It should be adjustable and unisex.

We named our solution Cogno-vest (See Fig. 3(b)). Cogno-vest is a Y-harness brace with stretchable straps, wrapped around the shoulder, chest, and lower back, securing garment positioning on the users’ torso. To support the wearable hardware, the back part of the brace, taken from the Mil-Tec Military-style Lightweight vest, covers the entire back. This piece of fabric is made of durable polyester nylon and integrated laser-cut loops, which facilitate mounting solenoid actuators on the back (see Fig. 3(d)). As a result, Cogno-vest is unisex, lightweight (the overall weight, including actuators and controller board, is 1 kg), and allows for unimpeded, free breathing. The stretchable material and Velcro fasteners make it size adjustable and body-conform.

#### 3) Solenoid boxes

The actuators were placed in custom 3D printed boxes so that they could be fixed to the wearable garment. These mechanically distance-adjustable, 3D printed boxes (see Fig. 3(c)) were designed to account for the irregular surface of the human back, especially on the spinal cord and lower back areas. The experimenter can manually adjust the distance of each solenoid with the participants’ back to ensure that there is contact between the tactile display and the user’s back. A 20 mm silicone tube was used (hardness: 60 shore A, inner diameter:12 mm, outer diameter: 14 mm) to extend the solenoid actuator tips, mainly, for actuators located on the spinal cord (actuator 4-6) and lower back areas, where higher curvature can be found, depending on the participants’ back-morphology.

#### 4) System Evaluation

In order to compare the performance of the integrated solenoid actuators to that of their datasheet, we tested one solenoid actuator under various environmental and parametric conditions, as shown in Fig. 4(a): *Solenoid* (upper figure: only solenoid), *Solenoid-Tip* (middle figure: solenoid with elastic tip), and *Solenoid-Box* (lower figure: only solenoid inside the 3D printed box). In each condition, we sent a square pulse signal (pulse width = 250 ms, pulse duration = 5.25 s) to the solenoid. We measured the provided impact force, activation delay (ActD: the time interval between command sending and solenoid activation), and deactivation delay (DeactD: the time interval between command sending and solenoid deactivation) at different stroke lengths of 0, 1, 2, 3, 4, and 5 mm (end of the stroke).

**Fig. 4.**
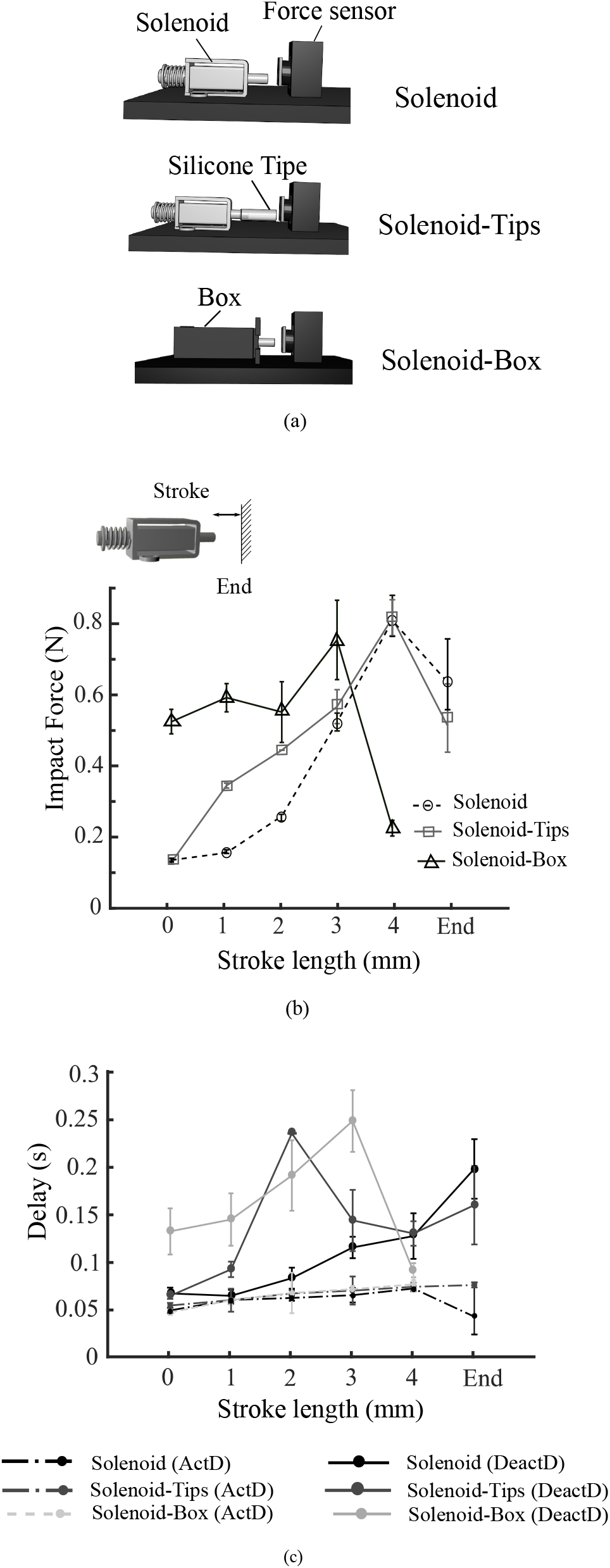
Solenoid characterization experimental setup and results. (a) Schematic view of solenoid characterization setup in three different experimental arrangements. (b) Provided impact force by solenoid at different stroke length in three experimental conditions. (c) Solenoid activation and deactivation delays as a function of stroke length in three experimental conditions.

The solenoid’s impact force profile, ActD, and DeactD are shown in Fig. 4(a) and Fig. 4(b). Fig. 4(b), indicates a gradual increase in the impact force profile with increasing stroke length for all three conditions. This observation suggests that in order to get a stronger touch sensation, each solenoid’s position needs to be adjusted to (approximately) 60-80% of the maximum stroke length. Moreover, the new arrangements did not reduce the impact force, although its dependency to the stroke length changed.

We also observed that delayed activation of 50 ms followed by a longer, variable deactivation delay, even for the *Solenoid* condition (Fig. 4(c)). While ActD is constant (around 50 ms), the deactivation delays increase with stroke length.

To conclude, the new arrangement of solenoid improves the force profile; however, it leads to bigger delays to follow the deactivation command. To compensate ActD and DeactD, we deployed a linear estimation that calculates front robot position for the next 100 ms (approximate mean of ActD and DeactD) with the current velocity and position.

## III OWN BODY PERCEPTION EXPERIMENT

### A. Participants

We recruited 25 healthy participants (12 female), aged between 18 and 32 years (mean age: 26±3.9 years). Participants were all right-handed, reported no previous neurological or psychiatric conditions, and were naïve to the purpose of the study. All participants gave written informed consent before participating and the research was conducted in accordance with the Helsinki Declaration.

### B. Experimental Design

We designed a behavioral study to investigate the feasibility of inducing PH in healthy participants by providing tactile-motor signals from the prototyped Cogno-vest. As in our previous studies [31], [33], we explored the effect of synchrony (synchronous (SYNC) vs. asynchronous (ASYNC)) in providing self-generated tactile stimuli on the participant’s back. There was no force feedback at the hand for both SYNC and ASYNC modes. To reduce the habituation effect and related potential changes in tactile perception of our participants, a variable delay of 500 ± 100 ms was used in the asynchronous block.

We used a 4-item bodily illusion questionnaire to estimate subjective changes in altered own body perception during the experiment. The questionnaire was adapted from [31] (experiment 2) and given to participants at the end of each experiment block. The statements used in the questionnaire can be seen below:

***Self-touch***—”I felt as if I was touching my back with my finger.”
***Passivity***—”It was as if someone else was touching my back.”
***PH***—”I felt as if someone was standing close to me or behind me.”
***Control***—”It was as if I had two bodies.”

We asked participants to report the degree of their agreement on a 7-point Likert scale for each item (0 = not at all, 6= very strong). In addition, we asked all participants to write down a short description (at least two lines) about any observation that they may have had during the experiment.

### C. Procedure

Participants wore the Cogno-vest over a fitted T-shirt. They were blind-folded and received noise-canceling headphones (WH-1000XM3, Sony) to eliminate the sound of solenoid activation during the experiment. The experiment consisted of two main blocks (see Fig. 5), namely the calibration task and the poking task.

**Fig. 5.**
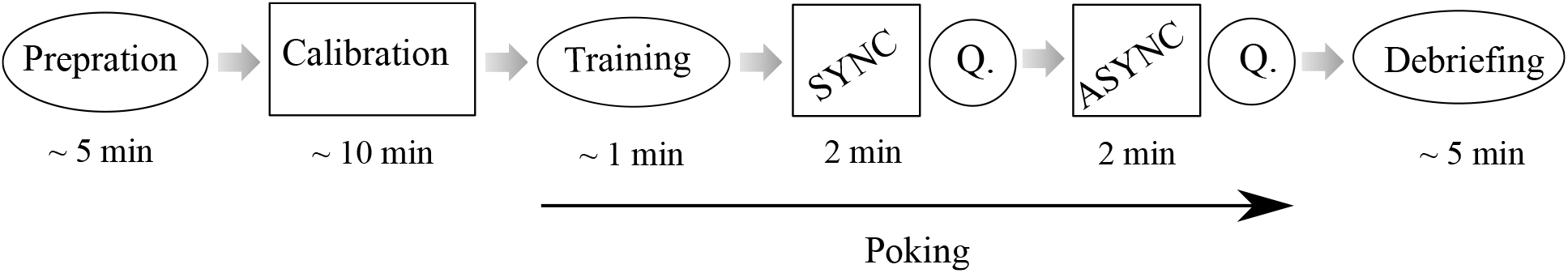
Experiment flow. In the preparation session, the experimenter helped participants to properly wear the Cogno-vest on a thin T-shirt. Then she manually calibrated the position of each single solenoid, ensuring that there is actual contact between solenoids and participant’s back (Calibration). Subsequently, participants started to complete two blocks of poking task (followed with bodily illusion questionnaire), after getting familiarized with the task. At the end of the poking task, participants commented on their experience and the setup freely (Debriefing).

#### 1) Calibration

In the calibration task, the experimenter activated each solenoid actuator individually (in a random sequence) to ensure that there was contact with the back. The experimenter manually adjusted the solenoid position at different stroke lengths until receiving verbal confirmation from participants that they felt mechanical touch from each solenoid. The calibration session lasted around 10 minutes. Participants were asked to stand during the whole experiment to avoid changing solenoid position.

#### 2) Poking task

Following [31] (experiment 2), we asked participants to produce poking-like hand movements with the front robot in order to they receive mechanical touch on their back (synchronously or asynchronously) through the Cogno-vest (see Movie S2).

The experiment started with a training session in which participants performed the poking task (synchronous), first with their eyes open, and then closed. We instructed participants to place their right-hand index finger inside the finger placement and generate forward and backward hand movements with the front robot at a frequency of approximately 1 Hz, while also instructing them to freely explore the entire workspace (up-down, left-right). The training session lasted around 2 minutes. During the subsequent poking task, participants were sound-isolated and blindfolded. In this way, they completed two blocks of SYNC and ASYNC (one block of each), equally distributed among participants. Each block lasted 2 minutes, and an acoustic cue was presented at the beginning and the end of each block. After each block, participants were asked to complete the questionnaire and, at the end of the poking task, to comment freely on their experience and the experiment.

### D. Results

The questionnaire data of the Cogno-vest experiment were collected from 25 participants. As the questionnaire data were not normally distributed (Shapiro-Wilk test for normality), we analyzed the effect of synchrony (Sync vs. Async) with the one-tailed Wilcoxon signed-rank test for each questionnaire item. The mean values for Self-touch, Passivity, PH, and Control questions in both SYNC and ASYNC conditions are presented in Fig. 6(a). We report first, that the average ratings for all three experimental items (*Self-Touch*, *Passivity*, *PH*), in both conditions, are significantly higher than those of the *Control* question.

**Fig. 6.**
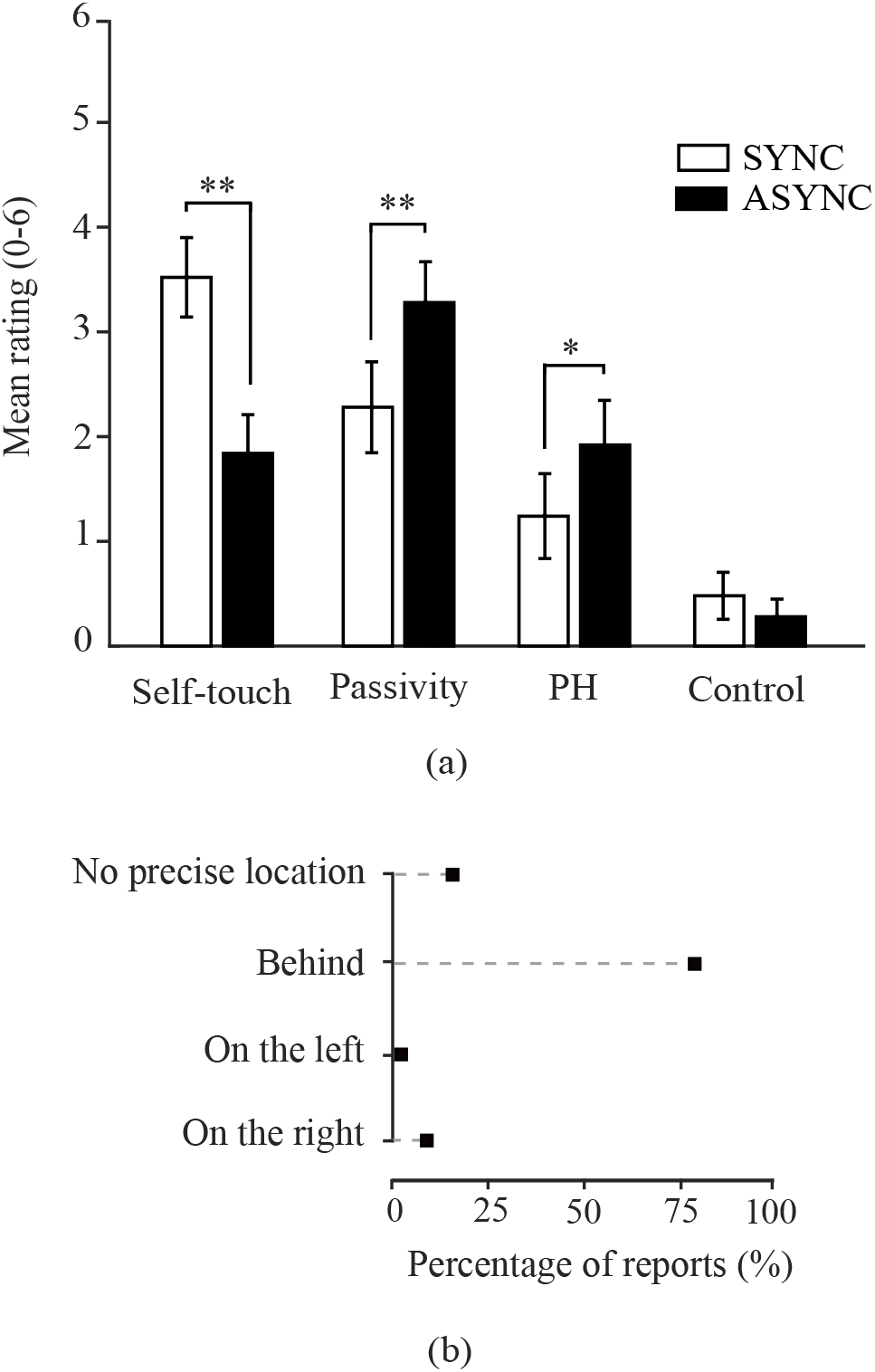
Results of bodily illusions questionnaire. (a) Mean ratings for bodily illusions and control questionnaire items. The bar charts and error bars depict the average scores and standard error of the mean (SEM) for each question (**: p < 0.01, *: p < 0.05). (b) Participants’ reports on the location of presence in asynchronous condition.

Second, for all three experimental questions, we found a significant effect of synchrony *(Self-touch:* z=3.7, p < 0.01, *Passivity:* z=-2.5, p < 0.01, PH: z=-1.8, p = 0.03). As hypothesized based on [31], [33], participants gave higher ratings for the *Self-touch* question in the synchronous than asynchronous condition (SYNC: M=3.5, SEM= 0.4; ASYNC: M = 1.8, SEM = 0.4), while for *Passivity* experience and *PH* questions, their ratings were higher in the asynchronous condition (*Passivity*, SYNC: M= 2.3, SEM = 0.4; ASYNC: M=3.2, SEM=0.4; *PH*, SYNCH: M = 1.24, SEM = 0.4; ASYNC: M=1.9, SEM=0.4).

In addition, we asked participants who gave *PH* ratings >0to specify the location of the perceived presence. Fig. 6(b) represents the percentage of reports on the location of the perceived presence in asynchronous conditions, indicating that participants mainly perceived the presence behind their body.

## IV. Discussion

While the majority of prior studies have used vibrotactile stimulus for substituting collision-type stimuli on the users’ torso in virtual environments, employing force feedback actuators may enhance the level of immersion. However, compared to vibrotactile devices, force feedback interfaces demand higher considerations in the interface design and handling procedure, as their perceptions depend upon the actual physical contact with the skin. With higher morphological changes in the torso area, maintaining the physical contact with the skin of the torso is problematic. As a solution, we prototyped Cogno-vest, a unisex, body-conforming torso-worn force display. We further validate its performance in providing realistic collision sensations in a sensorimotor perception paradigm.

Our results demonstrate that the Cogno-vest can be used to experimentally induce illusory self-touch and passivity experiences as well as presence hallucinations (PH) of mild to moderate intensity in a healthy population. Despite the substantial inter- and intra-subject variability in torso morphology, the Cogno-vest was able to reliably provide mechanical touches on the back. To our knowledge, this is the first study to investigate the design, implementation, and application of an untethered, torso-worn force display.

Our results further replicate and extend prior findings [31] on the robotic induction of bodily illusions in a healthy population by providing a sensorimotor mismatch between the right hand and the back. In our previous study [33], we were able to induce illusory passivity experiences via a vibrotactile interface, but were not able to modulate self-touch and PH. However, the present force display allowed us to also modulate illusory self-touch and PH and compared to study [33], we have observed stronger passivity experiences reported by participants. These results are in line with prior studies [34], [41], which have shown the importance of rendering realistic force stimuli to simulate collisions or physical interactions in virtual environments.

### A. Study limitations

We note, however, that compared to the study by Blanke at al. [31], participants gave slightly lower ratings across all questionnaire items in both synchronous and asynchronous conditions. The observed decrease in illusory ratings may be explained by the forces that can be applied by the solenoids as seen in the evaluation test, which revealed that the maximum provided force is substantially smaller than the average poking force obtained from the human finger poking experiment. Indeed, providing such weaker touch sensations on the back, which has low spatial tactile acuity and density of touch receptors likely reduced the level of immersion and may lead to weaker induction and modulation of robot-induced illusory own body perceptions.

We speculate that smaller synchrony-dependent modulation might be due to additional temporal delays in solenoid activation and deactivation commands. In spite of implementing linear estimation to reduce ActD and DeactD, some participants still reported the feeling of time-delays at deactivation moments during the synchronous condition. Given that the present experiment aimed at investigating the effects of temporal mismatch in a somatosensory-motor stimulation paradigm between the hand and back, unexpected alterations in the temporal delay are likely to negatively influence participants’ responses. We argue that future applications may resolve these technical limitations by deploying a more reliable solenoid actuator, which can guarantee the time precision and ensure a sufficiently strong poking force.

## V. Conclusion and Future works

In this paper, we addressed the complexity of designing a body-conform force display by following an iterative, multidisciplinary design procedure to prototype Cogno-vest: a novel, torso-worn force display to provide mechanical touch cues on participants’back. Further, we evaluated the Cogno-vest’s potential for 1) providing touch cues and measuring tactile perception, but also 2) the induction and modulation of altered states of bodily self-consciousness.

Our findings demonstrate that the Cogno-vest is capable of successfully inducing PH and passivity experiences during asynchronous stimulation as well as self-touch during synchronous stimulation in a healthy population. Based on these findings we propose technological improvements, leading to the induction of higher illusory ratings. We are confident that the prototype Cogno-vest, described here, may pave the way for future user-friendly, torso-worn force display technology to provide personalized, realistic touch sensations in virtual environments.

We are currently employing the Cogno-vest in tactile spatial resolution tests to quantitatively evaluate its performance, while it is worn on the user’s body. We believe that the results of such research would provide more practical information on the design and handling of forcefeedback stimulators.

Future work should concentrate on the use of proportional force actuators, instead of On-Off actuators, to simulate approaching speed and variable force during poking movements and to create a more realistic poking sensation on the back. In addition, the handling procedure may be standardized by implementing automated distance-adjustable boxes to uniformly adjust solenoid distance from the body in a more precise and efficient manner.

Moreover, the front robot, not portable in the present study, can be replaced by currently existing motion-sensing technology in combination with a finger-based haptic stimulator. This would make the system fully portable and wearable, and allow us to perform experiments on tactile perception and BSC outside of the laboratory setting. With respect to clinical translation, this equates to placing the device also in the patient’s home or in clinics to facilitate the study of specific hallucinations such as the PH (or other bodily illusions) in ecological settings (i.e., close to daily life situations). It is also of interest to employ the portable Cogno-vest in a mobile setting, allowing us to investigate bodily illusions during human locomotion.

## Supporting information

Movie_S1_FrontRobot

Movie_S2_Synchronous_Sensorimotor_Stimulation

